# The loss of both pUL16 and pUL21 in HSV-1 infected cells abolishes cytoplasmic envelopment

**DOI:** 10.1101/2024.11.10.622843

**Authors:** Kellen Roddy, Peter Grzesik, Barbara Smith, Nathan Ko, Sanjay Vashee, Prashant J. Desai

**Author notes:** Correspondence (PJD) (PJD).

## Abstract

Previously, we had developed synthetic genomics methods to assemble an infectious clone of herpes simplex virus type-1 (HSV-1). To do this, the genome was assembled from 11 separate cloned fragments in yeast using transformation associated recombination. The eleven fragments or “parts” spanned the 152 kb genome and recombination was achieved because of the overlapping homologous sequences between each fragment. To demonstrate the robustness of this genome assembly method for reverse genetics, we engineered different mutations that were located in distant loci on the genome and built a collection of HSV-1 genomes that contained single and different combination of mutations in 5 conserved HSV-1 genes. The five genes: UL7, UL11, UL16, UL21 and UL51 encode virion structural proteins and have varied functions in the infected cell. Each is dispensable for virus replication in cell culture, however, combinatorial analysis of deletions in the five genes revealed “synthetic-lethality” of some of the genetic mutations. Thus, it was discovered that any virus that carried a UL21 mutation in addition to the other gene was unable to replicate in Vero cells. Replication was restored in a complementing cell line that provided pUL21 in trans. One particular combination (UL16-UL21) was of interest because the proteins encoded by these genes are known to physically interact and are constituents of the tegument structure. Furthermore, their roles in HSV-1 infected cells are unclear. Both are dispensable for HSV-1 replication, however, in HSV-2 their mutation results in nuclear retention of assembled capsids. We thus characterized these viruses that carry the single and double mutant. What we discovered is that in cells where both pUL16 and pUL21 are absent, cytoplasmic capsids were evident but did not mature into enveloped particles. The capsid particles isolated from these cells showed significantly lower levels of incorporation of both VP16 and pUL37 when compared to the wild-type capsids. These data now show that of the tegument proteins, like the essential pUL36, pUL37 and VP16; the complex of pUL16 and pUL21 should be considered as important mediators of cytoplasmic maturation of the particle.

## INTRODUCTION

Herpesvirus genomes have the coding capacity in excess of 100 genes. Many of these gene products have functions that are clearly defined as to their roles in virus replication. However, there are several gene products, many of them structural components of the virion, whose functions and activities in infected cells remain a mystery and have yet to be completely elucidated.

The herpes simplex virus type-1 (HSV-1) virion is comprised of four structural components: an icosahedral capsid, which encloses the viral DNA genome; an electron dense asymmetrically distributed material, which immediately surrounds the capsid and is termed the tegument; and an outer membrane or envelope, which encloses the tegument and capsid and in which are embedded the viral glycoproteins [1–4]. Capsid assembly and DNA packaging into icosahedral capsids are nuclear events. Subsequent nuclear exit and cytoplasmic envelopment, involve the participation of a large and diverse set of ∼50 proteins.

The tegument is one of the most complex and diverse structures of the virion both in terms of protein composition and the functions encoded by the constituents of this structure. The tegument is comprised of a dense protein network that maintains this structure even when devoid of the virus envelope or capsid [5]. The virus specified polypeptides that comprise this structure include those that function to activate transcription, shut off host protein synthesis, uncoat the virus genome, phosphorylate virus proteins and others whose functions are still poorly defined, reviewed in [4, 6–11]. The tegument displays a duality of functions in virus replication due to the role the proteins resident in this structure play both at early and at late times in infection. The tegument proteins have been classified as belonging to either the inner or outer layer of the tegument based on their close association with either the capsid (inner) or envelop (outer) [2, 12–16]. What has become increasingly evident is the importance of the tegument proteins in the maturation process of the enveloped virus. To date, three tegument proteins resident in the mature virion have been shown to have a deleterious and complete lethal effect on the maturation process. These are VP16 [17, 18], pUL36 (VP1/2) [19–22] and the product of the UL37 gene [20, 23, 24].

The studies presented here build on our recent experiments using the synthetic genomics assembly line to construct HSV-1 genomes carrying single, double, triple, quadruple and quintuplet mutations in different combinations (for the multiple mutations) of five genes encoding the tegument proteins pUL7, pUL51, pUL11, pUL16 and pUL21 [25]. This astonishing feat, to generate in parallel these mutant viruses, could only be done using this modular assembly method. Several studies have identified protein interactions between pUL7-pUL51 [26, 27] and between pUL11-pUL16-pUL21 [27–36] but other than single mutations, many have not been probed using multiple/combinatorial mutagenesis except for the UL7-UL51 gene pair [26, 37]. These proteins are conserved in all three of the herpesvirus families, yet are not essential, at least for HSV-1, in cell culture [26, 38–45]. They are most likely important for pathogenesis and spread of the virus *in vivo* as shown by analysis of some of the mutants in mouse model systems [46].

Our scientific premise is based on several lines of evidence that have demonstrated these proteins specify redundant functions because they are not required for virus replication in cell culture. We believe that we can uncover the nature of these “redundancies” using the novel synthetic genomics assembly line to discover “synthetic-lethals”. To this end, we generated multiple HSV-1 genomes carrying different combinations of deletions in these five genes. The outcome of this investigation revealed that any mutant virus that carried a combination which included a deletion in the UL21 gene always displayed a lethal phenotype [25]. We further investigated the combination of UL16 and UL21 mutations because these proteins have a documented history of physical interactions in the infected cell [28, 30, 31]. The viruses carrying single mutations in these genes replicated in non-permissive cells albeit poorly. The double mutant virus displayed significant impairment in virus replication. When this virus was examined further, it was evident that the virus assembled DNA filled capsids and these particles were able to exit the nucleus but failed to acquire the envelop in the cytoplasmic compartment. Further examination of the DNA-filled C capsids revealed a significant reduction in the capsid association of VP16 (outer tegument) and pUL37 (inner tegument) proteins. This indicates that the pUL16-pUL21 complex is required for incorporation of these essential tegument proteins, revealing the complex nature of how HSV-1 capsids mature into infectious particles.

## METHODS

### Cells and Viruses

Vero cells and transformed Vero cell lines (G5-9) were all grown in minimal essential medium (alpha medium – Gibco Invitrogen) supplemented with 10% fetal bovine serum (FBS – Gibco Invitrogen) and passaged as described previously [47]. G5-9 is a subclone of the original G5 cell line isolated by Stan Person in 1993 [48]. This cell line carries a genomic fragment that includes UL16 and UL21 and thus complements mutants that carry deletions in these two genes. All stocks of HSV-1 viruses were amplified as also described by Desai et al. [47].

### Antibodies

Antibodies to VP16 (LP1) were generated by Professor Tony Minson (University of Cambridge). This is a well characterized monoclonal antibody to this protein and has a significant citation record. Rabbit antibodies to pUL16 and pUL21 were made by John Wills (University of Pennsylvania, Hershey) and these have strong validation in the literature. Rabbit antibody to VP23 was generated by our lab using whole protein purified from capsid preparations and has demonstrated specificity [48]. Monoclonal antibody to gD (clone DL6) was generated by Dr. Cohen (University of Pennsylvania) and kindly provided to us by David Johnson (Oregon Health Sciences Center). This is a well-established antibody to gD. Antibody to pUL37 (rabbit polyclonal) was generated by Frank Jenkins (University of Pittsburgh). Mouse monoclonal antibody MCA406 which recognizes both VP21 and VP22a was purchased from Serotec Inc. GFP rabbit antibody (ab183734) was purchased from Abcam.

### Cre excision

For the Cre excision, we used 2.5 μg of the DNA in a 50 μl volume reaction and used Cre enzyme (2 units/μl) (NEB). This was incubated at 37°C for 30 minutes and then the enzyme heat-inactivated at 70°C for 10 minutes. The whole 50 μl reaction was transfected into Vero or G5-9 cells using X-tremeGENE transfection reagent (Sigma-Aldrich) using the protocol previously [25]. The transfection was harvested 3 days post and sonicated to generate an infected cell lysate. This was serially diluted and used to infect cells in 96 well trays. Single plaques isolated were amplified and checked by Phire Hot Start II polymerase (Invitrogen) PCR assays to check for excision as described previously [25, 49]. Positives were amplified further to generate high titer working stocks.

### Growth curves

Vero cells (5 x 10^5^) in 12 well trays were either infected at a multiplicity of infection (MOI) of 0.01 or 10 plaque forming units (PFU)/ml. The cells were harvested at 24, 48 and 72 hours post-infection for the low MOI infections or at 24 hours post-infection for the high MOI infections. Cells were freeze/thawed three times and virus progeny titered on G5-9 monolayers.

### Western blot analysis of infected cell lysates

Vero cells (5 x 10^5^) were infected at an MOI of 10 PFU/cell and harvested 24 hours post infection. Cell pellets were lysed in 2X Laemmli buffer and 10% of this sample was resolved using Nu-Page 4-12% Bis-Tris gels (Invitrogen) and transferred to nitrocellulose membranes using the iBlot2 system (Invitrogen) as described by Luitweiler *et al*. [50]. Rabbit antibodies to HSV antigens were used at a dilution of 1:500. Blots were processed using the enhanced chemiluminescence (ECL) kit (GE Healthcare) or Clarity chemiluminescence kit (Bio-Rad) according the manufacturer’s protocol and imaged using the iBright 1500 Imager (Invitrogen).

### Fluorescence light microscopy imaging

For confocal imaging, RPE-1 cells (5 x 10^5^) were seeded in a 4-well borosilicate glass bottom chamber slide (Lab-Tek). Cells were infected with each virus at a MOI of 10 PFU/cell and overlaid with FluroBrite DMEM (Thermo Fisher) supplemented with 1% FBS. 12 hours after infection, cells were imaged on a Zeiss LSM 510 confocal microscope using 63X objective.

### Transmission electron microscopy (TEM)

Vero cells (5 x 10^5^ cells) in 12 well tissue culture trays were infected at an MOI of 10 PFU/cell and processed for transmission electron microscopy (TEM) experiments [23]. Infected cells were processed 16 h post-infection. Samples were fixed in 2.5% glutaraldehyde, 3mM MgCl_2_, in 0.1 M sodium cacodylate buffer, pH 7.2 for overnight at 4°C. After buffer rinse, samples were postfixed in 1% osmium tetroxide in 0.1 M sodium cacodylate buffer (1 h) on ice in the dark. Following a DH_2_O rinse and en bloc staining in 0.75% uranyl acetate for three hours, samples were dehydrated in a graded series of ethanol and embedded in Eponate resin overnight at 60°C. Thin sections, 60 to 90 nm, were cut with a diamond knife on a Leica UltracutE ultramicrotome and picked up with 2x1 mm formvar coated copper slot grids. Grids were stained with 2% uranyl acetate (aq.) and 0.4% lead citrate before imaging on a Hitachi 7600 TEM at 80 kV equipped with an AMT XR80 CCD.

### Capsid purification and analysis of composition

Vero cells (20 X 10^6^) in 100 mm tissue culture dishes were infected at an MOI of 5 and harvested after 24 h. Capsids from infected cells were released by treating infected cell pellets with 2x CLB [48] followed by 30 sec sonication. Next, capsids were separated using rate-velocity sedimentation on a 20-50% sucrose gradient. Capsid bands were visualized using light-scattering, and the C-capsid band was harvested by side-puncture. The capsid fractions were TCA precipitated and the pellets resuspended in 2X Laemmli sample buffer. Proteins were separated by SDS-PAGE on a NuPage 4-12% Bis-Tris gradient gels and stained using SYPRO Ruby stain according to the manufacturer’s protocol (Thermo Fisher). Proteins from the same C-capsids preparations were again separated by SDS-PAGE and transferred to nitrocellulose membranes using the iBlot2 transfer machine. Membranes were processed for immunoblotting as described above. The primary antibodies used were rabbit R2421 αVP23 [48], mouse monoclonal LP1 for VP16 [51], rabbit 780 αUL37C [52], rabbit 74 αpUL16 [32], and rabbit 121 αpUL21 [31].

Quantitation of protein bands was performed using the iBright 1500 (Invitrogen). Bands were manually drawn and the values for Local Background Corrected Volume were calculated by the iBright software. For each set of C-Capsids, these values from the pUL37C and the VP16 bands were normalized to the Local Background Corrected Volume value of VP23 from the same sample. The normalized values for each virus protein were then compared and analyzed using GraphPad Prism 9 software.

## RESULTS

### Cre excision of the BAC-YCp sequence in the mutant viruses

Previously, we had observed that the HSV-1 strain KOS yeast assembled genome had problems with replication in Vero cells. This was judged to be due to the presence of the BAC-YCp sequence in the virus genome. Removal of the sequence resulted in wild-type kinetics of virus replication [25]. Because we have the vector sequence bracketed by *loxP* sites we performed Cre-excision on all of our assembled genomes in order to remove the BAC-YCp element. We used an *in vitro* Cre excision method which gave us an efficiency of approximately 70% and more recently almost 90% [49]. Single plaques were isolated following transfection of cells and screened using PCR assays for excision of the vector sequence. These plaques were used to amplify the virus to obtain a secondary stock and subsequently high titer working stocks. All these viruses encode a VP16-Venus fusion protein which enables one to visually follow virus replication (Fig. 1a). This fusion does not affect the ability of the virus to replicate [25]. We typically passage all the mutant viruses in G5-9 cells because of the complementing activity provided *in trans*. G5 cells were transformed with the EcoR1 G fragment (HSV-1 KOS nucleotides 29281:45511) and pSV2neo [48]. This fragment encodes genes UL16 to UL21. This cell line can complement mutants in UL16, UL17, UL18, UL19, UL20 and UL21. G5-9 is a subclone of G5 that displays better complementing activity. When the mutant viruses were plaqued on Vero cells, the Δ16 and Δ21 mutant viruses gave rise to small plaques on Vero monolayers. Plaques were not observed on Vero cells when the double mutant virus was plated, only single fluorescent foci were observed (Fig. 1b).

**Fig. 1.**
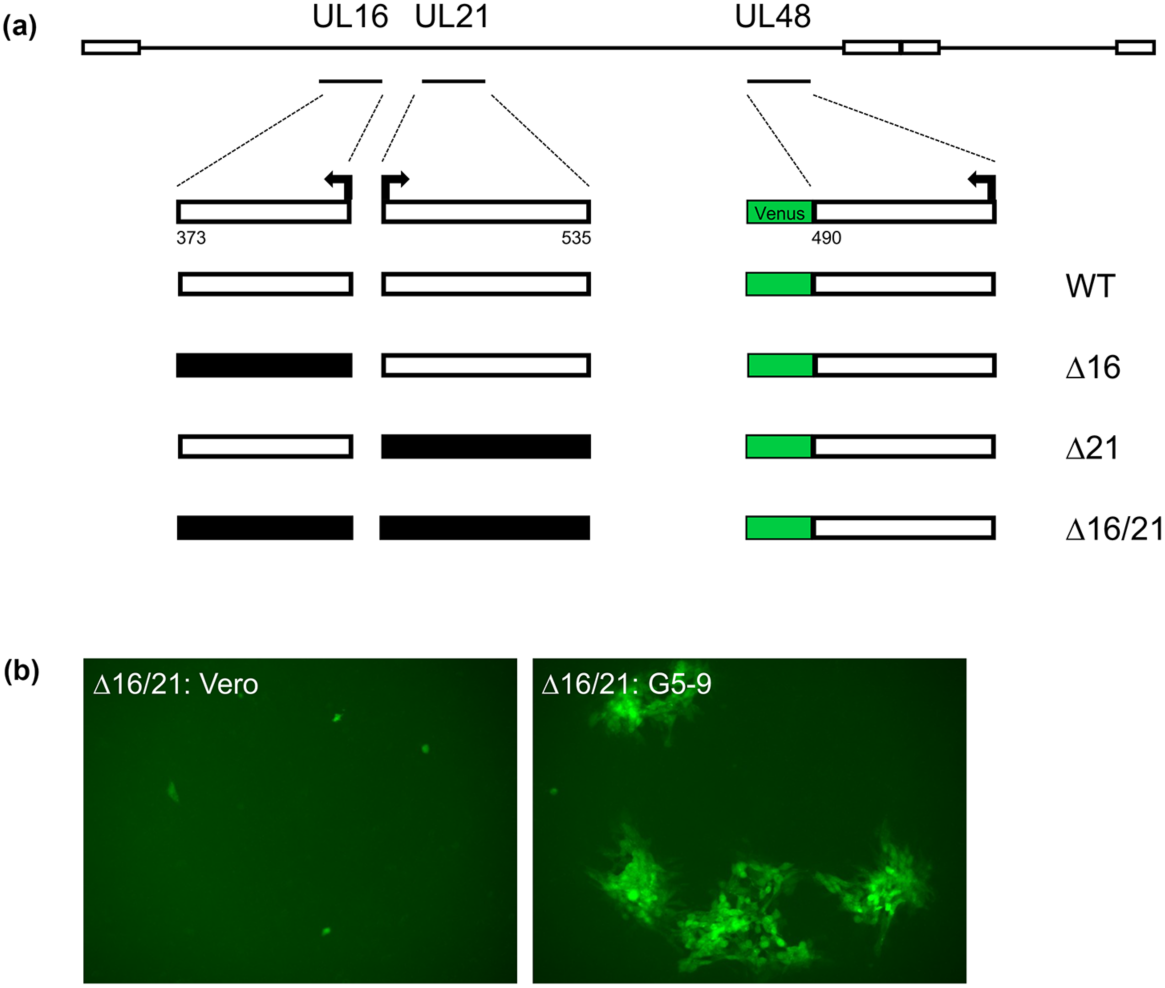
Illustration of the genotypes and phenotypes of the mutant viruses. (a) The genomes of the four viruses, WT (wild-type), Δ16, Δ21 and Δ16/21 (Δ:deletion) are shown. The deletions in the genes for UL16 and UL21 encompassed all the coding sequences. The Venus open reading frame (ORF) was fused to the C-terminus of VP16. Numbers of amino acid for each ORF are shown. (b) Fluorescence image of the plaquing efficiency of Δ16/21 on Vero and G5-9 cell lines (objective 10X).

### Protein expression

We used rabbit antiserum against pUL16 and pUL21 to confirm the expression of these proteins or their absence in cells infected with the different mutant viruses. Proteins pUL16 (predicted mass 40 kD) and pUL21 (predicted mass 58 kD) were observed in cells infected with the KOS^YA^ wild-type virus (Fig. 2). They were not observed in the cell lysates of the corresponding mutant and both proteins were absent in the double Δ16/21 mutant virus infected cells. We also examined the expression of other viral proteins in the same lysates. Thus, the levels of gD, VP22a, pUL37 and VP16 looked similar in both the wild-type and single mutant lysates. However, there was a detectable decrease in the levels of protein accumulation in the double mutant.

**Fig. 2.**
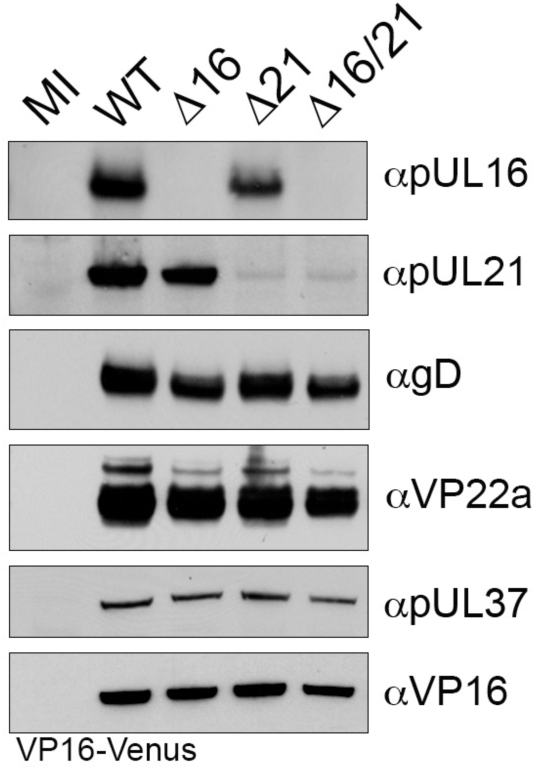
Expression of HSV-1 polypeptides in infected cells. Vero cells were synchronously infected with the indicated viruses or mock infected (MI). Cell lysates were collected 24 h post-infection and analyzed by SDS-PAGE and western blotting with antibodies for HSV-1 proteins. Both pUL16, or pUL21 are only observed in infected cell lysates for viruses expressing the corresponding wild-type gene. Additionally, lysates were probed with antibodies for gD, VP22a, pUL37, and VP16 which are all expressed with early and late gene kinetics and are unaltered in mutant virus lysates. Anti-VP16 antibodies detect the VP16-Venus fusion protein (92 kD).

### Growth curves

In order to quantitate the growth defect in the different mutants, we performed growth assays both at high multiplicity of infection (MOI) and at low MOI. Both the Δ16 and Δ21 mutant viruses could replicate on Vero cells but the yields of virus were lower than wild-type virus infected cells. At low MOI, there was a 2 log reduction in virus yield (Fig. 3a) and at high MOI there was a log reduction in virus titer (Fig. 4). For the double mutant there was negligible virus growth. The low levels of virus detected correspond to the amounts of input virus. This was also visually observed with the VP16 Venus fluorescence in low MOI infections. The Δ16/21 mutant was completely unable to spread to neighboring cells (Fig. 3b).

**Fig. 3.**
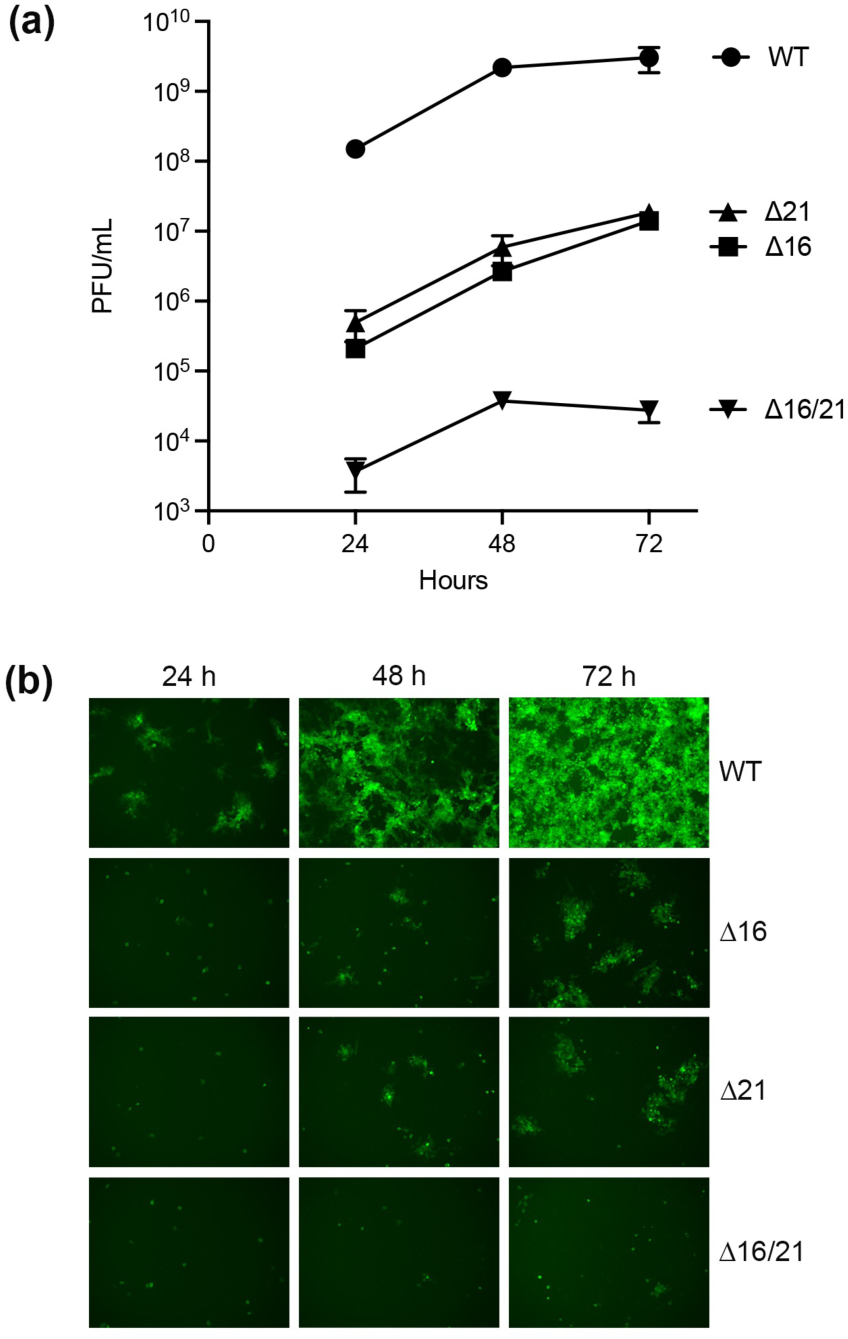
Viruses encoding deletions in UL16 and UL21 are attenuated for replication in Vero cells. (a) Vero cells were infected with each virus at a multiplicity of infection (MOI) of 0.01 plaque forming units (PFU)/cell and the infected cells harvested every 24 h over a 72 h period. Virus yield (PFU/mL) was enumerated by titration on G5-9 monolayers. Data from replicates was plotted for the multi-step growth curves. (b) Representative images of Vero cells infected at an MOI of 0.01 PFU/cell from the same cultures as above were obtained by fluorescence microscopy to visualize VP16-Venus expression at the different times post-infection.

**Fig. 4.**
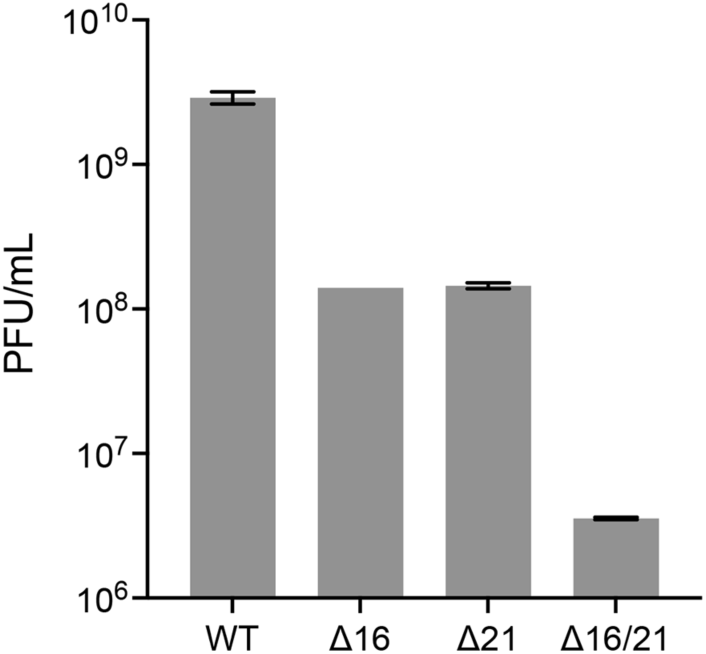
Single-step growth curves of the mutant viruses. Vero cells were infected at an MOI of 10 PFU/cell and the virus progeny harvested 24 h post-infection. Virus titers were enumerated by plaquing on G5-9 cells. Data presented are representative of two biological replicates.

### Confocal imaging of infected cells

Because the mutant viruses have the VP16-Venus tag in their genomes, we could visualize the fluorescence distribution using confocal light microscopy. For this, RPE-1 cells were used and infected at high MOI. Cells were imaged at 12h post-infection. VP16-Venus has a nuclear punctate distribution early in the infection but as time progresses, fluorescence is visualized at the nuclear and cytoplasmic membrane including the plasma membrane. In the cells infected with Δ16 mutant virus, the distribution of fluorescence was similar to wild-type. In the cells infected with Δ21 and Δ16/21 mutant viruses, the distribution of VP16 was perturbed and was less localized to the nuclear and cytoplasmic membranes. This was more evident for the double mutant virus (Fig. 5).

**Fig. 5.**
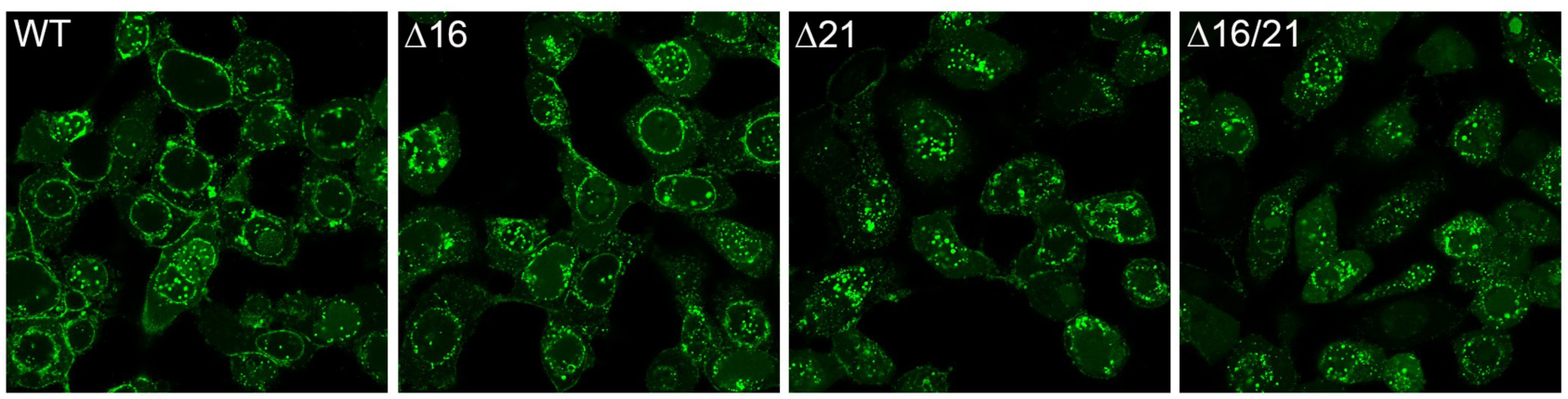
Confocal microscopy reveals disrupted perinuclear VP16-Venus localization in infected cells lacking pUL21. RPE-1 cells were plated in chamber slides and synchronously infected with each virus for 12 hours prior to live cell imaging by confocal fluorescence microscopy to visualize VP16-Venus (objective 63X). Perinuclear VP16-Venus was observed in WT virus and Δ16 infected cells, but this distribution became irregular in either the Δ21 or Δ16/21 infected cells.

### Ultrastructural analysis of infected cells

Vero cells were infected with all the mutant viruses and examined by electron-microscopy to visualize in greater detail what was happening within the cell (Fig. 6). For the wild-type infected cells, microscopy showed capsids in the nucleus, enveloped virus in the cytoplasm and at the cell surface, which is typical of productive virus production (Fig. 6a). In the cells infected with Δ16 (Fig. 6b) and Δ21 (Fig. 6c) mutant viruses, capsids were evident in the nucleus as well as in the cytoplasm. There were fewer enveloped viruses observed which reflects the lower production of virus in these cells. For the double mutant infected cells, we observed capsids in in the nucleus as well as in the cytoplasm (Fig. 6d). However, there were very few or no enveloped viruses detected in these cells.

**Fig. 6.**
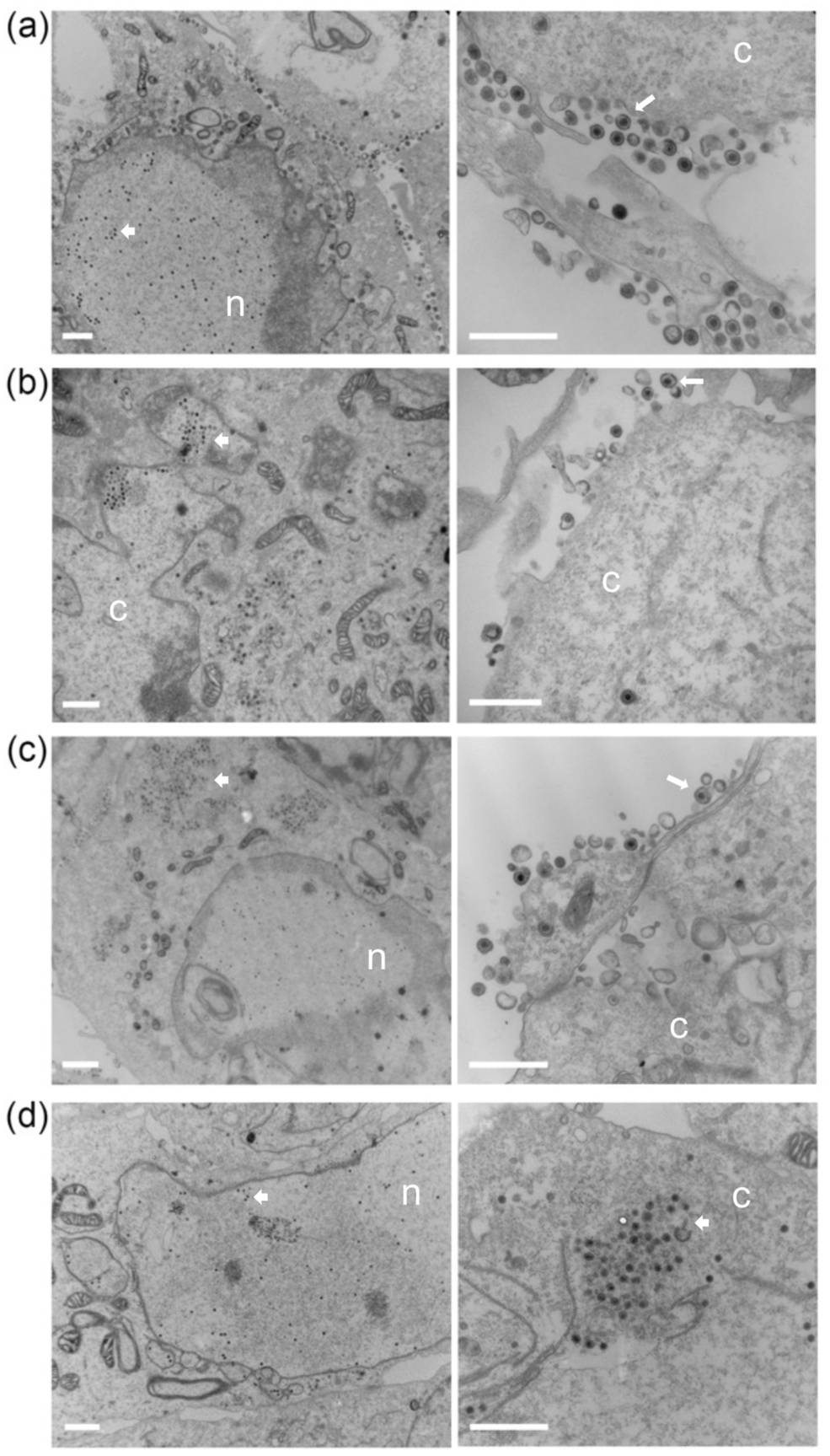
Transmission electron microscopy shows enveloped capsids in single Δ16 or Δ21 null virus infected cells, but not in Δ16/21 infected cells. Vero cells were synchronously infected with WT (a), Δ16 (b), Δ21 (c), or Δ16/21 (d) viruses (scale bar 1 µm) then fixed 16 h post-infection and processed for TEM imaging. Enveloped virus particles were observed in the cytoplasm and egressing from WT infected cells (white arrows) and observed at a lower frequency in Δ16 or Δ21 infected cells. Enveloped virus particles were not observed in Δ16/21 cells and an accumulation of unenveloped capsids (white arrowheads) was observed in the cytoplasm of these cells. The nucleus (n) and cytoplasm (c) are marked.

### Capsid assembly

We next examined capsid assembly and composition using purified capsids (Fig. 7). Whole cell lysates were sedimented through sucrose gradients and all three capsid types (A, B and C) were observed (Fig. 7a). There were lower levels of capsids in the gradients using lysates for Δ16/21 infected cells as judged by light scatter. We extracted the C capsids and analyzed these using total protein stain (Fig. 7b). One could readily identify the major capsid proteins, however, the levels of these and thus C capsids was generally also lower in the double mutant lysate gradients. We have analyzed a number of capsid gradients following replicate infections. The experiment shown in (Fig. 7b) shows lower levels of capsids in Δ16 virus cell lysates, which was not commonly seen. We next examined these C capsids for their composition using available antibodies to the different tegument and capsid proteins (Fig. 7c). We chose to normalize our capsids using antibody to VP23 which is present in capsids in a fixed amount (600 copies). When antibodies to VP16 and pUL37 were used, we observed a significant decrease in the amounts detected relative to VP23 normalization. This was examined using the quantitation software in the iBright 1500 and analyzed using GaphPad Prism9 software (Fig. 7d). There was a significant reduction in the capsid association of both VP16 and pUL37 in the mutant capsids. This same observation was observed in multiple replicate analyses of C-capsids isolated from infected cells using the same methods.

**Fig. 7.**
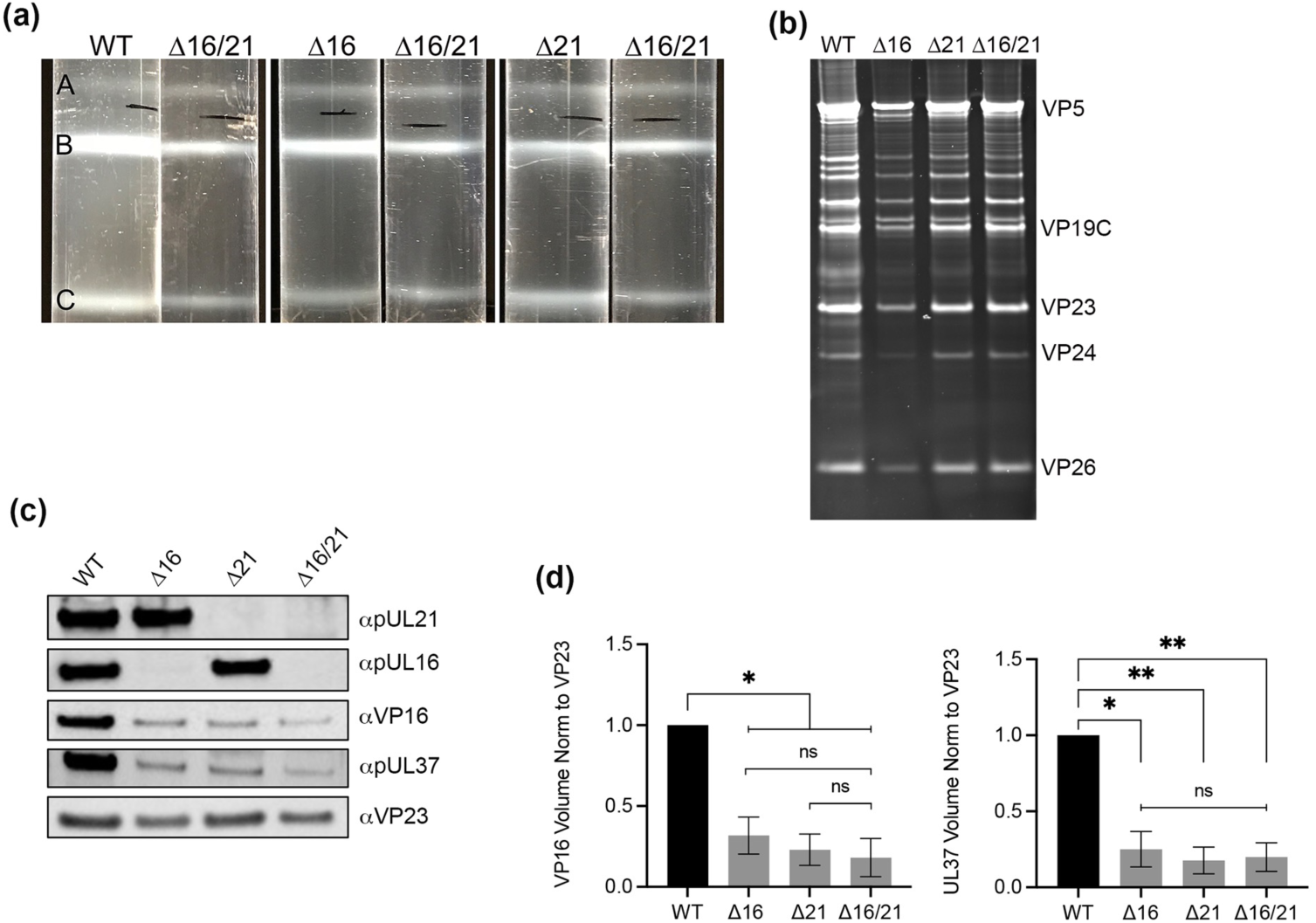
Isolation and analysis of mutant capsid particles. Vero cells were infected with each virus (MOI=5) and infected cell pellets were collected 24 h post-infection. Capsids from infected cells were released by treating infected cell pellets with 2X CLB and sonication followed by separation of capsids on a 20-50% sucrose gradients and ultracentrifugation. (a) Each capsid form — A, B, and C — are observed as light scattering bands and denoted on the gradient images, each compared to the Δ16/21 capsid bands. (b) C-capsids were harvested by side puncture from each gradient and proteins were separated by SDS-PAGE and observed by Sypro Ruby staining. Visible capsid protein identities are indicated. (c) Proteins from the same C-capsids were again separated by SDS-PAGE and probed for pUL16, or pUL21 by western blotting as well as the capsid triplex protein VP23. Additionally, capsid proteins were also probed for inner-tegument protein pUL37 or outer-tegument protein VP16 (Venus-fusion). (d) Quantitation of levels of VP16 and pUL37 detected in the C-capsid fractions relative to the triplex protein, VP23. Western blots were analyzed using the iBright 1500, yielding values of the Local Background Corrected Volume for each protein band. The VP16 and pUL37 volumes were normalized to the VP23 volumes, and then normalized to the WT capsids. Statistical analyses was performed with GraphPad Prism 9 using Student’s *t*-test. ns: not significant, *P*=<0.05 (*,**,***).

## DISCUSSION

Initial envelopment of the HSV-1 virion takes place at the inner nuclear membrane (INM). The interacting proteins, pUL31 and pUL34, the latter a membrane protein, are required for this initial envelopment; reviewed in [10, 11, 53–59] as well in some situations the US3 kinase. After the capsid is enveloped at the INM, it fuses with the outer nuclear membrane (ONM) depositing a naked (non-enveloped) particle into the cytoplasm [60]. These capsids are transported to the trans-Golgi compartment (TGN) or other cytoplasmic organelle (late endosomes) for final envelopment [10, 61–63]. This cytoplasmic site must accumulate all the different tegument proteins that are incorporated into the mature virion [4] and also the lipid membrane that envelopes this particle has to contain the full repertoire of viral glycoproteins, reviewed in [6, 10, 12, 56, 64–67]. One of the most intriguing aspects of this morphogenesis pathway is the role of the tegument proteins in this dual envelopment process, the cellular localization and movement of tegument proteins prior to their incorporation into the maturing virus and the viral factors/signals that traffic particles to the maturation compartment. What is still unclear is the composition of the tegument as the virus is translocated from the nucleus to the cell surface. The multitude of tegument proteins have different locations within the cell; some are exclusively cytoplasmic and others exclusively nuclear and yet others that are detected in both compartments. Thus, as the virus particle progresses on its way to the surface, all of these tegument components must be incorporated into the final mature virion. The mechanism by which it does this is still poorly understood. The manner by which protein-protein interactions determine the fate of virus particle formation is still unclear [6]. In fact the tegument proteins appear to be required for the transition of the capsids from the site of assembly to the cytoplasmic site for final envelopment. Regardless most observations demonstrate the tegument primarily matures in the cytoplasm and sequential interactions between capsid-tegument, tegument-tegument and tegument-envelope drive the assembly of this structure [29, 68]. One of the key questions is what complex synthesizes the tegument assembly. Data has been interpreted that suggests [69–71] it is the largest tegument protein, pUL36 that initiates this. Data has also demonstrated the role of VP16 in complex with VP22, pUL41 and pUL47 as the “organizers” of the tegument [29, 70, 72–74]. Data has also implicated the complex comprised of pUL11, pUL16 and pUL21 playing a pivotal role in linking capsids with the envelope [28, 30–32, 35, 36, 75].

Clearly three tegument proteins, pUL36, pUL37 and VP16 are absolutely required for secondary envelopment. Additional gene products of the virus are also required, however, redundancy of activities in tissue culture may hide their distinct roles in this process. Proteins, pUL16 and pUL21 have been identified in different activities in the infected cell. The UL21 gene product has RNA binding activity, it is involved in syncytial processes, affects US3 kinase activity, inhibits innate immunity signaling, acts as a viral phosphatase and alters host metabolic pathways [76–82]. Protein UL16 similarly displays a variety of activities in the cell including interactions with host mitochondria [83, 84] as well as a role in syncytial formation [85]. Both pUL16 and pUL21 exhibit dynamic interactions with both tegument and glycoproteins of the virus [6, 28, 29, 32–35, 43, 85–88]. Deletion of each gene individually has been done in a number of HSV strains with differing results [40, 41, 43, 89–93]. In most cases, deletion of the gene in HSV-1 does not significantly affect virus replication, albeit it can impact the levels of virus production. In HSV-2, the single deletion of UL16 or UL21, affects nuclear egress and retention of the viral genome in the assembled capsid [91, 94, 95]. Recent studies also demonstrate that a mutant with deletions in both UL16 and UL21 fails to dock to the nuclear pore [96]. These different activities illustrate the complexity of the functions of these two proteins in HSV infected cells. In our study in KOS HSV-1 strain, we do not see nuclear egress defects as judged by ultrastructural analyses but this could be due to strain differences between this family of viruses [93].

In our study, it is evident that pUL21 as well as pUL16 mediate important interactions that affect capsid association of tegument proteins. This includes the inner-tegument protein pUL37 and a major constituent of the tegument, VP16. The reduction of tegument protein incorporation affects robust virus replication (as in the case of the single deletions) or completely abolishes virus envelopment (as in the case of the double deletions). Both these proteins have extensive interactions with tegument and envelop proteins. pUL16 has been shown to interact with VP16 as well as gE, pUL11 and VP22 [28, 32–35, 43] and is required for the virion incorporation of gD [87]. Thus, the pUL16-pUL21 complex could be required for bridging interactions between the inner and outer teguments and subsequently with the envelop during secondary envelopment.

## Author Contributions

KR, PG, BS, NK, SV and PD carried out the experiments. KR, PG, BS, NK, SV and PD wrote the manuscript and generated the figures.

## Acknowledgements

We want to thank John Wills (Penn State Medical School) for antibodies to pUL16 and pUL21. Also, Professor Tony Minson (University of Cambridge) generously provided the LP1 monoclonal for our use. Finally, David Johnson (Oregon Health Sciences Center) and Frank Jenkins (University of Pittsburgh) for anti-gD and anti-pUL37 antibodies, respectively.

